# Fine-Grained Structural Classification of Biosynthetic Gene Cluster-Encoded Products

**DOI:** 10.64898/2025.12.12.693975

**Authors:** Vladimir Porokhin, Emily Mevers, Justin J. J. van der Hooft, Soha Hassoun

## Abstract

Biosynthetic gene clusters (BGCs) are responsible the biosynthesis of many natural products, including a multitude of effective therapeutics and their precursors. Advances in genomic data collection as well as computational techniques have made it possible to identify BGCs at scale. However, accurately determining the types of BGC-encoded products from genomic content remains elusive. Here, we introduce *BGCat* (BGC annotation tool), a machine learning method for fine-grained structural classification of BGC-encoded products, leveraging the NPClassifier natural product nomenclature. Our method leverages a pre-trained protein language model for creating meaningful gene representations and a deep neural network for class label prediction. We show the method outperforms state-of-the-art approaches in coarse-grained product classification and is effective for detailed classification. We implement a clustering-based augmentation strategy for BGC-product relationships, addressing a crucial gap in the available datasets. We then introduce the concept of product class profiles (PCPs) of gene cluster families (GCFs), associating each GCF with a probabilisitc distribution of product types and offering a new perspective on GCF functions. Lastly, we use *BGCat* to provide new product class labels for over 100k BGCs in antiSMASH DB that presently have minimal information about their products.

## 1 Introduction

Natural products have been a staple of medicine for much of human history. Natural products represent a vast reservoir of potential drug candidates, with a high proportion of biologically-active species [1, 2]. A large percentage of drugs in current use owe their existence to natural products, including antibiotic lariocidin [3], anti-tumor agent anthramycin [4], antiparasitic drug Ivermectin [5], and approximately 70% of anti-infective drugs [6]. By fall of 2019, over a third (38%) of FDA-approved new molecular entities were either natural products or their derivatives, and in select areas (e.g. cancer drugs) that percentage can be as high as 65% [1]. As such, there is an ongoing interest in natural products and their sources.

In microbes, the specialized genes that are used to synthesize natural products are clustered into biosynthetic gene clusters (BGCs). These genes encode enzymes that synthesize and tailor the structure of the product, as well as regulatory, and transport proteins involved in its synthesis [7]. The recent explosion of microbial genomic data has led to the development of computational techniques for identifying BGCs. Early methods relied on sequence alignment algorithms and manually curated rules. However, with the rapidly growing quantity of identified BGCs and their diversity, algorithmic and rule-based approaches have given the way to machine learning techniques, e.g. DeepBGC [8], BiGCARP [9], and BGCCGB [10], promising a more flexible and comprehensive framework for understanding BGCs. AntiSMASH (ANTIbiotics and Secondary Metabolite Analysis SHell), currently the most widely used tool for mining microbial genomes for BGCs [11], leverages profile Hidden Markov Models in conjunction with a bank of rules for detecting and characterizing BGCs, with the latest version supporting up to 101 cluster types [12].

However, despite advances in BGC detection, determining the identities of the natural products encoded by BGCs remains challenging. For example, although antiSMASH infers the approximate scaffold for some metabolites based on PKS (polyketide synthease) and NRPS (non-ribosomal peptide synthease) domains and includes some analysis features for tailoring enzymes [12], it currently does not predict the full product structure with tailoring modifications. Several methods were proposed for product structure prediction, including RiPPMiner [13], TransATor [14], and PRISM [15]; however, they are either limited to specific types of BGCs or are unable to recognize novel enzymatic activities, restricting their general applicability. Recently, CHAMOIS was introduced as a generalized model for characterizing chemical properties of BGC products based on Pfam domains [16]; however, its ClassyFire-based [17] classification system was primarily designed for the wider bio-organic chemistry community and as such is less than ideal for natural products [18]. Meanwhile, the traditional approach to BGC product classification, as seen in DeepBGC [8], BiGCARP [9], and BGCCGB [10], employs a coarse-grained set of classes provided in MIBiG (Minimum Information about a Biosynthetic Gene cluster) [19]. This nomenclature currently categorizes products into 7 classes: Alkaloids, Polyketides, NRPs (non-ribosomal peptides), RiPPs (ribosomally-synthesized and post-translationally modified peptides), Saccharides, Terpenes, and Others. Because these approaches rely on broad, coarse product categories, they offer limited insight into the structural diversity of the encoded metabolites. This gap highlights the need for fine-grained structural classification methods capable of distinguishing natural products at a resolution meaningful for biosynthetic and chemical analysis.

The lack of high-quality data associating BGCs and natural products has been a major obstacle for gleaning insight into product structures. For example, the largest repository of such information, MIBiG, provides a selection of 3,013 BGCs [19]. Consequently, a common strategy has been to group BGCs into gene cluster families (GCFs), which can link BGCs to better-annotated MIBiG entries that include structural information as a starting point for BGC characterization [20, 21]. Implementations of this approach typically rely on network analysis [22], sequence similarity [23, 24], or a combination thereof [25, 21], with statistical correlation techniques sometimes introduced to leverage mass spectrometry data [23, 26]. Grouping clusters in such fashion provides important context for understanding BGCs and has been shown to create meaningful families that are comparable to chemical classes [21]. However, the focus thus far has been on establishing connections to known clusters as opposed to learning more general patterns of BGC product expression.

We introduce in this paper *BGCat* (BGC annotation tool), a technique for fine-grained structural classification of BGC-encoded natural products following the NPClassifier nomenclature designed for natural products [18]. Our technique leverages ESM Cambrian, a protein language model trained on a wide range of protein sequences [27], as the engine for condensing relevant biosynthetic genes into meaningful embeddings. We show that our approach surpasses the state-of-the-art techniques for traditional coarse-grained BGC product classification. Next, we demonstrate that our technique is effective for detailed product classification. We then introduce a clustering-based augmentation method that provides additional data for model training and enhances its performance. Next, we investigate product class profiles (PCPs) of GCFs to help identify the most meaningful families. Finally, we leverage *BGCat* to introduce detailed product labels for thousands of BGCs in the antiSMASH database (antiSMASH DB) [28], including the many clusters with unknown products. We make our model available at https://github.com/HassounLab/BGCat.

## 2 Methods

### 2.1 Dataset construction

Our dataset is derived from MIBiG, the largest manually curated source of experimentally validated BGCs along with their products. The database is supported by an online community of 250+ members and currently includes 3,013 clusters [19], 96% of which are associated with bacteria and fungi. To use the most reliable BGC annotations, we exclude 377 of those clusters that are marked as retired, resulting in a set of 2,636 MIBiG BGCs. Next, we leverage NPClassifier [18] to obtain detailed product type labels for these BGCs. The nomenclature used by NPClassifier is designed to be informative for natural products and features a hierarchy with three distinct levels: 7 pathways, 70 superclasses, and 672 classes; however, not all of them are covered by the BGC products. The product structures are processed as SMILES strings, which we canonicalize and de-duplicate using RDKit [29] to ensure each unique product is only classified once. When a BGC is known to produce multiple products, their labels are combined in a multi-label fashion. This process results in a dataset associating BGCs and their corresponding product classifications. In the set of MIBiG BGCs, 521 clusters lacked machine-readable product structures (e.g. a valid SMILES string was not specified) and an additional 69 BGCs produced no NPClassifier predictions; both of these BGC types were excluded from further consideration. This process yielded 2,046 BGCs, featuring 3,479 product structures, together resulting in 3,839 unique BGC-product pairs that formed our core MIBiG dataset. The most common compound categories at the pathway level were Polyketides and Amino acids & Peptides, with 77% and 65% of the BGCs featuring at least one product of the respective type. On average, each product was associated with 1.15 unique pathway labels.

### 2.2 Model architecture and training

We structured our model as a feed-forward deep neural network. The network operates on BGC gene embeddings and predicts product classification in a multi-label fashion. We implemented three separate networks for each of the pathway, superclass, and class label types, as well as a combined network that predicts all three types simultaneously. The BGC embeddings are constructed by applying the ESM Cambrian 600M [27] model to all biosynthetic (core and additional) genes; the remaining gene types (regulatory, transport-related, and other) were excluded due to their incidental effect on the product structure. Mean pooling was then used to combine gene-level embeddings into whole-BGC embeddings. The input dimension of the network is 1,152 to match the size of the BGC embeddings, the hidden layers progressively reduce in their dimensionality, and the final layer predicts the target labels. The model consists of three hidden layers implemented using PyTorch and is trained using the Adam optimizer [30] with binary cross-entropy as the loss function. The architecture of the model is illustrated in Figure 1.

**Figure 1.**
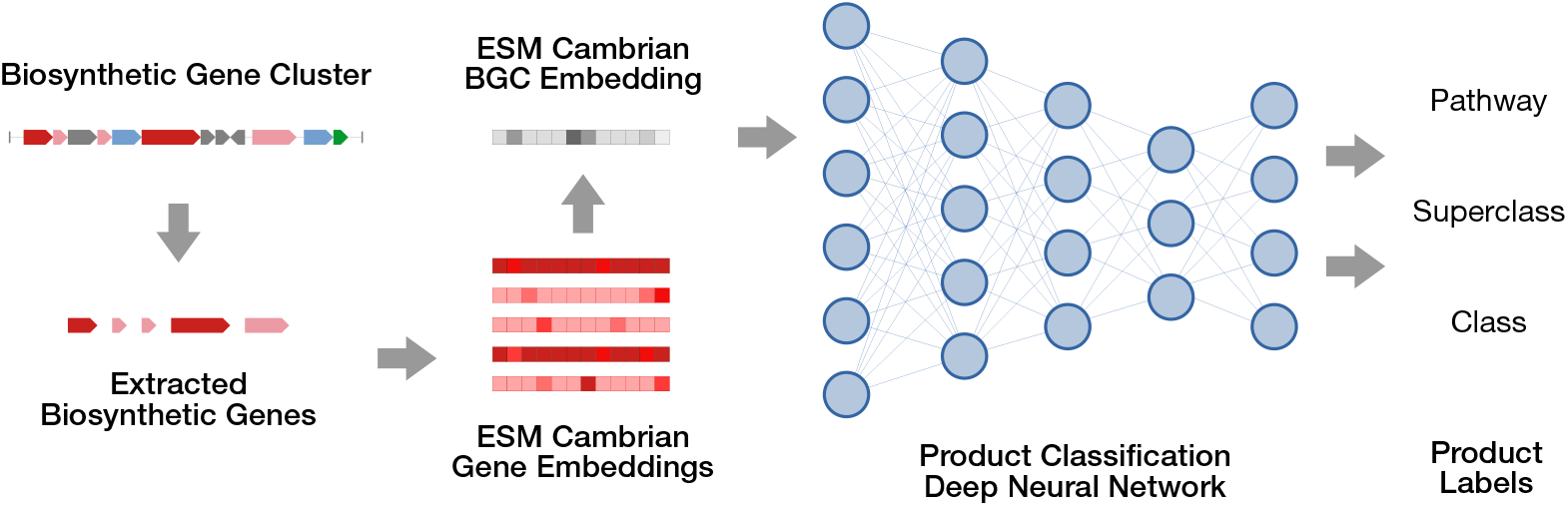
An overview of the *BGCat* model. A BGC identified by antiSMASH or a similar tool is accepted as the input. Next biosynthetic genes, shown as shades of red, are extracted and embedded using ESM Cambrian. Regulatory, transport-related, and other genes, represented by green, blue, and gray arrows, respectively, are ignored. The gene-level embeddings are then aggregated to yield an embedding representative of the BGC. A deep neural network is then used to predict pathway, superclass, and class labels of the potential products.

### 2.3 Dataset augmentation

The core MIBiG dataset was augmented with BGCs sourced from BGC Atlas [31], an online database containing nearly 1.8 million clusters, derived from 35k bacterial metagenomes found in MGnify [32]. These BGCs were identified by antiSMASH and as such include both complete and incomplete clusters. Incomplete BGCs pose a problem in this instance as they may be insufficient to support product classification; we therefore focused on the subset of 204,661 complete clusters. In addition, the BGC Atlas clusters provide limited details about their product molecules, so we sought to leverage the product information available in similar BGCs in MIBiG. Following the method described in BGC Atlas, we grouped the complete BGC Atlas clusters into GCFs using BiG-SLiCE 2.0.0 [21] with a distance threshold of 0.4, resulting in a set of 18,596 GCFs. Next, we queried our core MIBiG BGCs against those GCFs, finding at least one similar MIBiG BGC for 743 GCFs from the set. In the event BiG-SLiCE identified multiple candidate GCFs for a given BGC, only the highest-ranking assignment was used. We assume that all BGCs within a given GCF will yield substantially similar products; therefore, product classification labels from the MIBiG BGCs were copied to all BGC Atlas-derived BGCs that are part of the same GCF. This process resulted in a set of 9,055 BGCs sourced from BGC Atlas with their corresponding product labels. These BGCs, combined with our core MIBiG set, resulted in 11,101 BGC-label pairs which we refer to as our augmented set.

## 3 Results

### 3.1 Method outperforms existing techniques in traditional BGC product classification

Currently there is no comparable approach that seeks to classify BGC products based on a detailed nomenclature appropriate for natural products. Most of existing work in this space treats BGC product classification as an incidental problem to BGC detection, classifying products into the 7 coarse-grained classes provided in MIBiG [19]. Although this nomenclature provides somewhat limited information about the product, it offers a benchmark where we can compare against current baseline techniques. We considered four methods as our baselines: the built-in classification functionality of antiSMASH [12], DeepBGC [8], BiGCARP [9], and BGCCGB [10]. We evaluated the model on two versions of the MIBiG database: version 1.4 to allow for direct comparison with previous work, and version 4.0 to use the most up-to-date information. In both cases, the evaluation was performed using 5-fold cross-validation.

As shown in Table 1, our approach provides substantial performance gains over the baselines in traditional BGC product classification. In a head-to-head comparison using the MIBiG v. 1.4 database, our method improved upon AUROC and precision relative to prior approaches, although the recall was lower than that of BGCCGB. However, this gap in recall disappeared when the model was evaluated on MIBiG v. 4.0. These trends help illustrate two key areas where our method has improved over existing techniques. First, the use of ESM Cambrian embeddings of gene sequences alongside a deep neural network resulted in substantial gains over all but one baseline methods. Second, the introduction of additional BGC information available in a more recent release in MIBiG allowed our model to better learn the product classification task and surpass all baselines.

**Table 1.** Average AUROC, recall, and precision for traditional BGC product classification; table adapted from BGCCGB[10].

| Metric | antiSMASH | DeepBGC | BiGCARP | BGCCGB | Our method with |  |
| --- | --- | --- | --- | --- | --- | --- |
|  |  |  |  |  | MIBiG 1.4 | MIBiG 4.0 |
| AUROC | 0.784 | 0.792 | 0.855 | 0.862 | 0.936 | **0.937** |
| Recall | 0.718 | 0.691 | 0.539 | 0.740 | 0.693 | **0.768** |
| Precision | 0.658 | 0.741 | 0.593 | 0.732 | 0.811 | **0.825** |

### 3.2 *BGCat* is effective for BGC natural product class prediction

Next, we apply our method for detailed natural product class prediction. In this problem, we consider the three NPClassifier classification levels with increasing level of specificity: pathway, superclass, and class [18]. The pathway level consists of seven classes: Alkaloids, Amino acids and Peptides, Carbohydrates, Fatty acids, Polyketides, Shikimates and Phenylpropanoids, and Terpenoids. As such, pathway-level classification is similar to MIBiG class prediction seen in traditional BGC product classification, although the mapping between the two is not one-to-one. Meanwhile, the set of possible superclasses and classes is substantially larger, with an almost order of magnitude increase at each level. We consider two types of models: one that predicts a specific level of classification, and another that predicts all levels simultaneously.

To provide a comprehensive evaluation of the method, we consider four metrics: average precision (AP), area under the receiver operating characteristic (AUROC), recall, and precision. The latter two provide an approximate measure of performance assuming a fixed likelihood threshold of 0.5, which is a customary default choice [33] for binary classifiers (e.g. whether or not a BGC makes a product of a particular class). However, such a cut-off is not necessarily optimal and its selection is subject to trade-offs [33, 34]. Therefore, we also consider AUROC, a traditional classifier performance metric, as well as average precision, a measure of the quality of relative ranking. A key characteristic of our BGC dataset is class imbalance, with some types of BGC products being vastly overrepresented compared to others. We therefore employ two averaging schemes for our metrics. Micro averaging treats each prediction in a standalone fashion, representing the performance of a typical prediction. Macro averaging computes the metric for each class separately and reports the average across the classes. With all classes being treated equally irrespective of any imbalance, macro averaging represents the performance of the method for a randomly selected class. When applied to average precision, this macro-averaged value corresponds to the mean average precision (mAP) commonly used in multi-label classification.

The results calculated with 5-fold cross-validation on the MIBiG 4.0 set can be found in Table 2a. The method achieves high AUROC values across the board; however, other measures are more varied. In general, performance reduces with increasingly more specific classification levels. The prediction of all levels simultaneously presents intermediate performance. Recall values are considerably lower with class-level predictions, suggesting that many classes cannot be identified, likely as a consequence of class imbalance. This is further supported by the significantly lower macro averaging results, where the underrepresented classes have greater impact on the metric. However, precision remains high, indicating that the model is relatively accurate in the predictions it makes.

**Table 2.**
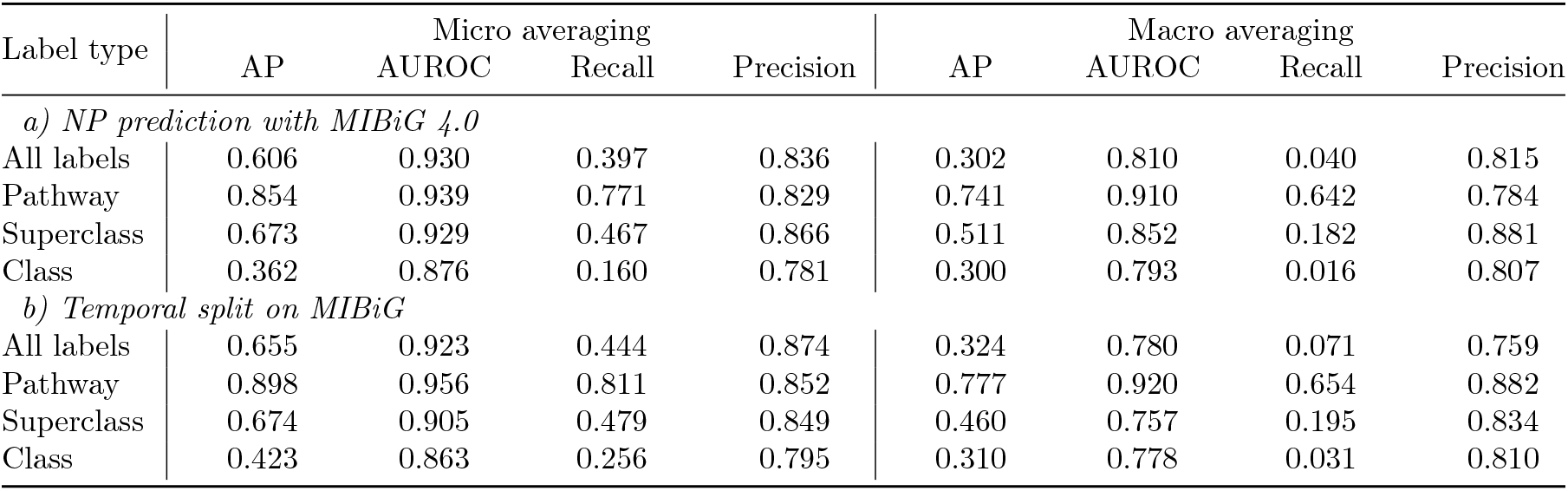
NP label prediction performance, for all labels being predicted at the same time and pathway, superclass, and class labels being predicted separately. a) Results for a random 5-fold cross-validation split. b) Results for a temporal split on MIBiG, with updates between versions 3.1 and 4.0 used as the test set.

| Label type | Micro averaging |  |  |  | Macro averaging |  |  |  |
| --- | --- | --- | --- | --- | --- | --- | --- | --- |
|  | AP | AUROC | Recall | Precision | AP | AUROC | Recall | Precision |
| *a) NP prediction with MIBiG 4.0* |  |  |  |  |  |  |  |  |
| All labels | 0.606 | 0.930 | 0.397 | 0.836 | 0.302 | 0.810 | 0.040 | 0.815 |
| Pathway | 0.854 | 0.939 | 0.771 | 0.829 | 0.741 | 0.910 | 0.642 | 0.784 |
| Superclass | 0.673 | 0.929 | 0.467 | 0.866 | 0.511 | 0.852 | 0.182 | 0.881 |
| Class | 0.362 | 0.876 | 0.160 | 0.781 | 0.300 | 0.793 | 0.016 | 0.807 |
| *b) Temporal split on MIBiG* |  |  |  |  |  |  |  |  |
| All labels | 0.655 | 0.923 | 0.444 | 0.874 | 0.324 | 0.780 | 0.071 | 0.759 |
| Pathway | 0.898 | 0.956 | 0.811 | 0.852 | 0.777 | 0.920 | 0.654 | 0.882 |
| Superclass | 0.674 | 0.905 | 0.479 | 0.849 | 0.460 | 0.757 | 0.195 | 0.834 |
| Class | 0.423 | 0.863 | 0.256 | 0.795 | 0.310 | 0.778 | 0.031 | 0.810 |

To showcase the performance of the method as additional BGC information becomes available, we introduce a temporal data split. In this instance, we retained BGCs in a previous version of MIBiG (v. 3.1) for training the model, and reserved the BGCs added in the next version (v. 4.0) for testing the performance. The results can be found in Table 2b. In this case, the performance of the method is broadly similar to that with random cross-validated split; however, some metrics are varied likely due to the smaller test set size: the test size in the cross-validated split was 768, while in the temporal split it was 183. Nevertheless, this indicates the method remains effective on newly added BGCs.

### 3.3 Dataset augmentation enhances BGC NP class predictions

The core MIBiG dataset is restrictive for machine learning methods that require large amounts of data. As such, we sought to increase the number of training examples by introducing a data augmentation scheme that has expanded the dataset size 5-fold. The effect of that change on prediction performance can be seen in Table 3a. The evaluation was performed on the same temporal split as in Table 2b, and the BGCs reserved for testing were excluded from the augmentation scheme. We note a consistent increase in all metrics except for precision. This indicates the model was able to recall more of the classes with enhanced quality of ranking, at the expense of slightly lower accuracy of its predictions.

**Table 3.** NP label prediction performance with dataset augmentation. a) Results for a temporal split on MIBiG, with updates between versions 3.1 and 4.0 used as the test set. b) Results for the augmentation split, where the augmented examples were used for training and non-augmented ones for testing. c) Results for the realistic data split, with larger GCFs used for training and smaller unseen GCFs for testing.

| Label type | Micro averaging |  |  |  | Macro averaging |  |  |  |
| --- | --- | --- | --- | --- | --- | --- | --- | --- |
|  | AP | AUROC | Recall | Precision | AP | AUROC | Recall | Precision |
| *a) Temporal split* |  |  |  |  |  |  |  |  |
| All labels | 0.669 | 0.937 | 0.554 | 0.784 | 0.426 | 0.854 | 0.235 | 0.750 |
| Pathway | 0.889 | 0.962 | 0.837 | 0.809 | 0.760 | 0.936 | 0.689 | 0.783 |
| Superclass | 0.637 | 0.924 | 0.521 | 0.731 | 0.436 | 0.856 | 0.310 | 0.601 |
| Class | 0.495 | 0.896 | 0.413 | 0.735 | 0.361 | 0.835 | 0.196 | 0.796 |
| *b) Augmentation split* |  |  |  |  |  |  |  |  |
| All labels | 0.442 | 0.933 | 0.508 | 0.404 | 0.135 | 0.819 | 0.078 | 0.238 |
| Pathway | 0.596 | 0.871 | 0.742 | 0.481 | 0.453 | 0.824 | 0.599 | 0.408 |
| Superclass | 0.403 | 0.915 | 0.459 | 0.337 | 0.177 | 0.832 | 0.144 | 0.248 |
| Class | 0.201 | 0.894 | 0.270 | 0.312 | 0.124 | 0.810 | 0.054 | 0.300 |
| *c) Realistic split* |  |  |  |  |  |  |  |  |
| All labels | 0.315 | 0.835 | 0.271 | 0.531 | 0.112 | 0.646 | 0.049 | 0.455 |
| Pathway | 0.527 | 0.790 | 0.505 | 0.530 | 0.403 | 0.734 | 0.367 | 0.443 |
| Superclass | 0.304 | 0.807 | 0.242 | 0.562 | 0.129 | 0.691 | 0.067 | 0.386 |
| Class | 0.151 | 0.772 | 0.127 | 0.448 | 0.094 | 0.628 | 0.032 | 0.452 |

Next, we validated whether the BGCs introduced by augmentation were informative by introducing an augmentation split, where the clusters sourced from BGC Atlas were used for training the model and the remaining ground-truth MIBiG BGCs were reserved for testing. The results can be found in Table 3b. With lack of access to high-quality MIBiG BGCs, the performance of the method has significantly decreased; however, it remained effective to a degree, suggesting that the added BGCs provide valuable information for the product classification task.

### 3.4 Model generalizes to unseen GCFs

With the landscape of known and well-annotated BGCs being limited, it is highly desirable for the model to be applicable to novel BGCs with no previously described function. To estimate the performance of the method under those circumstances, we introduce a “realistically novel” data split similar to one described in Profile-QSAR [35]. This split seeks to maximize the difference between training and test sets by separating BGCs according to their GCF membership. The training set is assembled from the BGCs comprising the largest GCFs, while the smallest GCFs are used to build the test set. This reflects a degree of realism because the largest GCFs are likely to contain more common BGCs with a greater likelihood of appearing in our dataset. Conversely, the likely less common BGCs in the smallest GCF are reflective of novel BGCs that we might not encounter in the course of training our model.

The results of this experiment can be found in Table 3c. Compared to the temporal split in Table 3a, the performance is considerably lower; however, the model retains a degree of effectiveness on the unseen GCFs, demonstrating its generalizability.

### 3.5 BGC product classification offers a new perspective on GCFs

Detailed classification of BGC natural products enables more comprehensive understanding of GCFs. Existing methods for constructing GCFs primarily leverage genomic sequence information for grouping similar BGCs, and although they do not directly consider the natural products, the resulting families are often correlated with specific product structures [22, 23]. However, it remains uncertain whether this pattern holds universally for all types of products and BGCs. As such, there is value in considering the natural products more explicitly.

To offer greater insight into GCFs, we propose the concept of product class profiles (PCPs), whereby each GCF is associated with a probability distribution of natural product types. We construct these distributions accordingly, by applying our model to these BGCs and aggregating the predicted types for each GCF. Thus, a given GCF may be associated with specific classes of molecules to varying degrees. Of particular interest are the GCFs with a distinct PCP, whose product type distribution is meaningfully different from the overall distribution of the dataset (the background), as such GCFs are likely to be strongly associated with particular types. We identify these GCFs using Pearson’s chi-squared test with a significance threshold of p *<* 0.05. This process yielded 5,083 (27%) distinct GCFs out of the 18,596 derived from BGC Atlas, and we report these GCFs in Supplementary Tables 1 and 2.

Select examples of such PCPs are shown in Figure 2. GCF 942 (Figure 2a) is representative of an indistinct GCF, whose product type distribution is not significantly different from the background. The two most common pathways, Amino acids & Peptides and Alkaloids are represented in this family in a similar proportion to the overall dataset. In contrast, GCF 778 (Figure 2b) has a distinct distribution with a rich variety of product types, with classes from the Carbohydrates, Polyketides, and Shikimates & Phenylpropanoids pathways notably overrepresented. GCF 11048 (Figure 2c) is also an example of a distinct GCF, albeit a more specialized one for production of carbohydrates.

**Figure 2.**
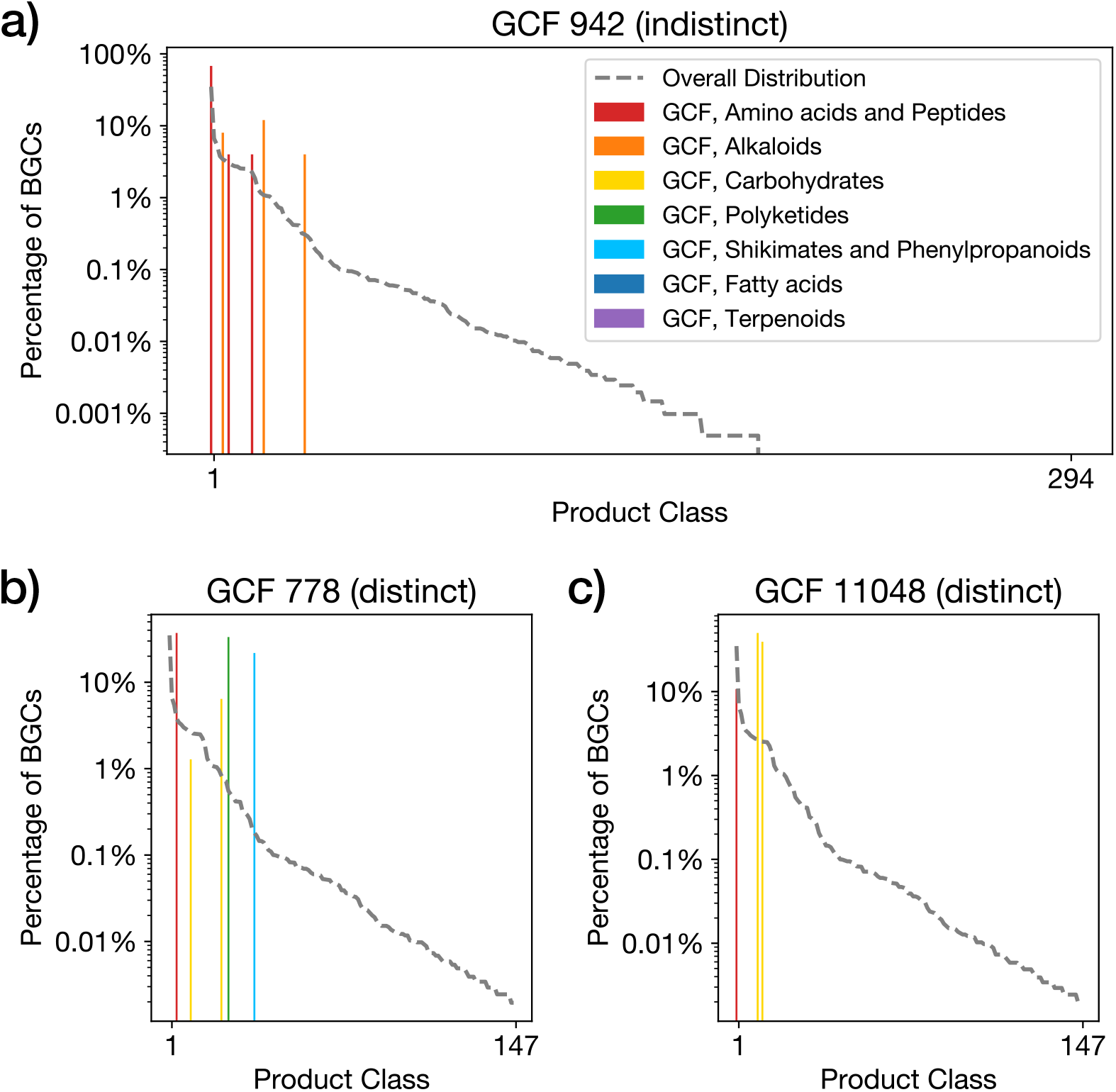
Example product class profiles for different GCFs constructed using *BGCat*. (a) A GCF with an indistinct distribution of product types: amino acids and alkaloids are represented similarly to the overall dataset. (b) An example of a GCF with a distinct distribution of product classes featuring a multitude of product types. (c) A GCF with a distinct distribution specialized in specific product classes. In both distinct cases, the proportion of BGCs responsible for a given product type is well above the average for the dataset (dashed gray line).

### 3.6 *BGCat* predicts detailed product labels for microbial BGCs in anti-SMASH DB

To showcase a practical application of our method, we apply it to antiSMASH DB [28], a public repository of over 260k BGCs from over 36k bacterial genomes precomputed using antiSMASH. Although antiSMASH is able to elucidate the approximate scaffold of some products, in most cases it is unable to predict the full structure or detailed classification of the product. One way to gather deeper context about the product is to leverage the KnownClusterBlast (KCB) module of antiSMASH that identifies similar clusters in MIBiG and thus can provide structures of related natural products. However, in some cases, the MIBiG BGCs may not include the product structure, or a KCB match may not be available for a particular antiSMASH BGC.

We can therefore separate antiSMASH DB into three subsets. Subset 1 consists of 114k BGCs with both a KCB match and one or more product structures available in the corresponding MIBiG BGC. Subset 2 contains 27k BGCs that have a KCB match but no machine readable structure provided in MIBiG. Subset 3 includes 121k BGCs that feature no KCB matches at all. The first two subsets offer an opportunity to evaluate *BGCat*, since the MIBiG BGCs identified by KCB will contain a coarse classification of the product at minimum; subset 1 additionally provides product structures, which can be used to compare NPClassifier’s performance to the same standard. The final subset offers opportunity for new discovery using our prediction approach. An overview of the three subsets can be found in Supplementary Figure 1.

To evaluate our model, we implement a mapping from NPClassifier terms to MIBiG coarse human-annotated labels and consider the percentage of BGCs for which at least one of *BGCat* predictions is in agreement with the MIBiG classification. This measure effectively represents our model’s ability to recall information from MIBiG BGC labels. These labels, being annotated by human experts, can be considered highly accurate; however, due to individual preferences or unknown BGC products these labels may not necessarily be exhaustive. Most categories map trivially; however, Shikimates and Phenylpropanoids had no direct equivalent and had to be mapped to the MIBiG “Other” category. Fatty acids were mapped to Polyketides in MIBiG due to the close structural and evolutionary similarities between their corresponding synthases [36]. RiPPs and NRPs are difficult to distinguish based on the peptide structure alone, so they were grouped into one surrogate MIBiG label “RiPP/NRP.” The objective of the model is therefore to predict a detailed product class that maps to the same label found in a MIBiG KCB match.

On subset 1, the top prediction of the model is able to recover a correct MIBiG label for 76% of the BGCs; the percentage increases to 84% when top two predictions are considered. NPClassifier on average also predicts two labels per BGC, and its rate of agreement with MIBiG is similarly at 85%. Because our dataset is based on the product classification from NPClassifier, this result represents the upper bound on the *BGCat* ‘s accuracy and its ability to nearly match it is encouraging. On subset 2, the machine readable structure of the MIBiG product is not available, which precludes the use of NPClassifier. However, our model is still able to recover at least one MIBiG label for 54% of BGCs with its top prediction, and for 67% of BGCs with its top two predictions. The lower performance is expected in this case because the lack of structure in a MIBiG BGC likely means the structure is not part of the MIBiG database, which would prevent our model from observing it in the training set.

Finally, we consider subset 3, for which no KCB data is available. In this case, *BGCat* can provide unique insights about the products of these BGCs. We offer our top 3 predictions for all of the 121k BGCs as the Supplementary Table 3. To evaluate utility of these new results relative to the available GCF groupings, we calculated the Normalized Mutual Information (NMI) between the GCF assignments and the labels. NMI is an established metric for evaluating the agreement between two clusterings based on information theory. For pathway-, superclass-, and class-level labels, the NMI was found to be 0.16, 0.27, and 0.38, respectively. These low values indicate that product labels and GCF membership share relatively little mutual information and therefore capture largely orthogonal aspects of BGC biology: GCFs reflect sequence similarity, whereas *BGCat* ‘s predictions reflect chemical outcomes. The increasing NMI from pathway to class level further suggests that finer-grained product labels align more closely, but still only modestly, with genomic similarity, whereas coarse labels obscure these relationships. Thus, *BGCat* provides substantial, nonredundant information beyond what GCF groupings alone can offer.

Figure 3 illustrates the breadth and diversity of product types predicted across antiSMASH DB subset 3. At the pathway level (Figure 3a), all seven natural product pathways are well represented, with Amino acids & Peptides and Polyketides dominating both BGC and GCF counts. This broad coverage suggests that *BGCat* predictions span the full metabolic landscape of microbial natural products. In contrast, the class-level distributions (Figure 3b) exhibit a characteristic long-tail pattern: a handful of classes contain thousands of BGCs, whereas many others are sparsely populated. This imbalance underscores both the chemical richness of BGC-encoded metabolites and the difficulty of fine-grained classification. Notably, GCF counts broadly track BGC frequencies, indicating that highly populated classes correspond to genuinely diverse biosynthetic families rather than duplicated clusters. In other words, classes with many BGCs also contain many distinct genomic architectures, showing that their abundance reflects true biosynthetic diversity rather than repeated instances of the same cluster. Together, these figures validate the utility of fine-grained structural classification and highlight the data challenges inherent to learning at this level of resolution.

**Figure 3.**
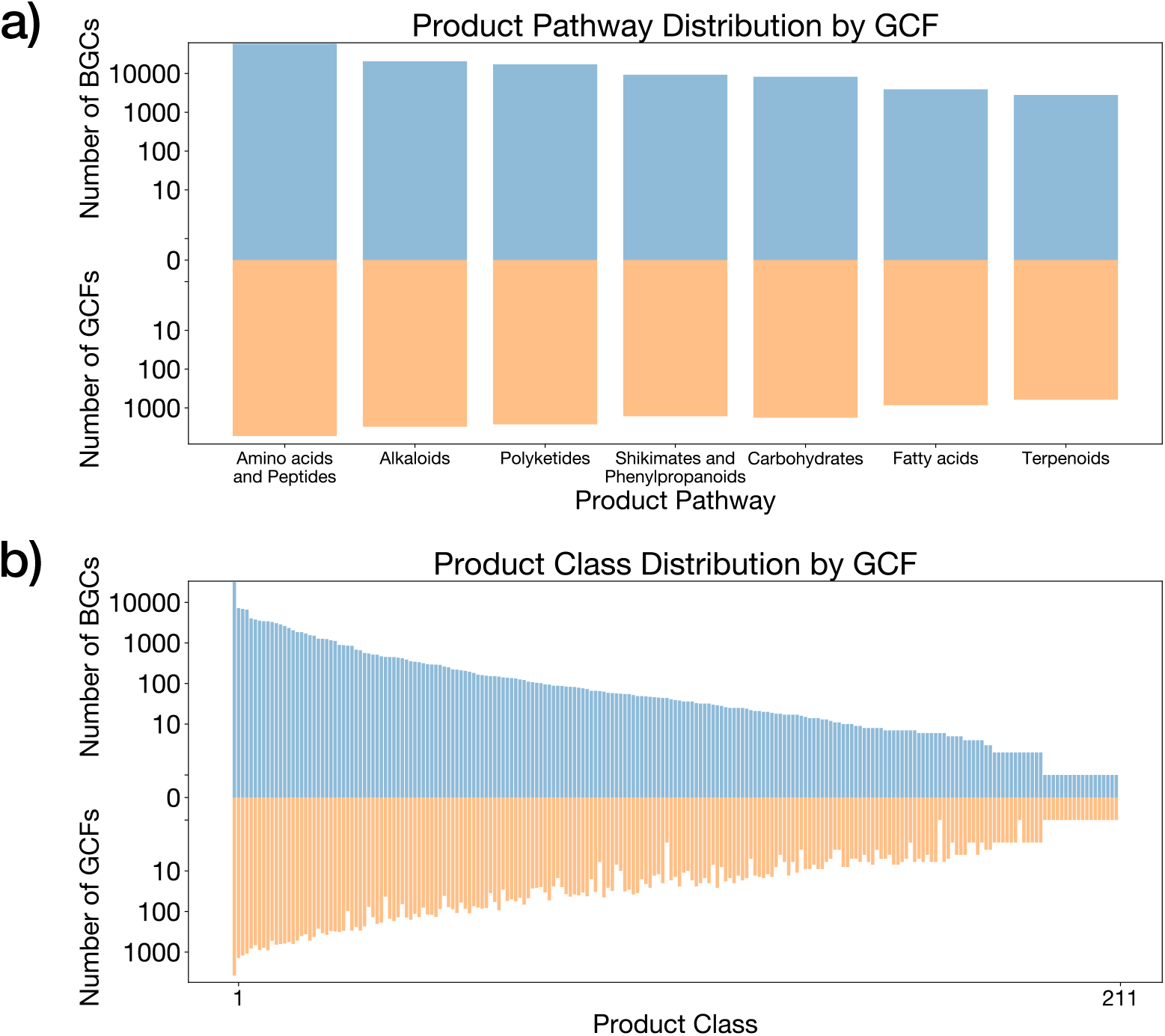
Distributions of BGCs and GCFs in the antiSMASH DB subset 3, stratified by: (a) the 7 pathway labels, and (b) product class labels. The x-axis represents different product labels predicted by the model. The y-axis represents the number of BGCs or GCFs which contain a given product class. The BGCs are shown as blue bars in the upper half of each plot, while the GCFs are shown as orange bars in the bottom half. These distributions highlight the broad biochemical coverage of subset 3 and reveal substantial fine-grained diversity, underscoring the value of *BGCat*’s product-based predictions for characterizing BGCs beyond sequence-based groupings.

## 4 Discussion

*BGCat* is a deep neural network for fine-grained BGC product classification. It builds upon the NPClassifier [18] nomenclature for natural products, making it particularly suitable for describing metabolites arising from BGCs. By leveraging a large pre-trained protein language model, our method takes advantage of the vast repository of available protein sequence information to create biologically meaningful representations of biosynthetic genes. In addition, our novel data augmentation strategy allows the neural network to learn from a greater variety of BGCs than would otherwise be possible with presently existing datasets. We show that our method is effective both for traditional coarse-grained as well as fine-grained BGC product classification. The fine-grained approach is especially interesting in the context of BGC products as there is currently no method capable of predicting the complete structure of such molecules in the general case; thus, detailed product classification offers the greatest insight into the function of BGCs currently available. Furthermore, we anticipate that *BGCat* can improve paired omics analyses by improved linking and ranking of predicted BGCs in genomes to mass spectra with predicted compound classes [37].

In contrast to prior algorithmic methods, our machine learning-based approach aims to be generalizable to different types of clusters and products. We have shown that the model is effective for predicting product classes for new BGCs introduced over time and maintains a degree of accuracy on novel types of BGCs when grouped by GCFs. As new information about BGCs becomes available, our model can be trained on new data to further enhance its accuracy and coverage of product types. It should be noted, however, that our model may need further adaptations to apply beyond microbial BGCs, as BGCs found in other organisms often require special considerations [38, 39, 40].

To highlight a practical application of our method, we have used it to provide product labels for a diverse set of microbial BGCs deposited in antiSMASH DB. We have shown that our model provides product class labels in agreement with MIBiG labels for at least 54% of BGCs with KCB matches. However, of particular interest was the subset of 121k BGCs with no KCB matches: in this case, our model provided the only means of predicting the BGC product type.

While our tool was effective in different BGC product classification scenarios, it has a few limitations that could be addressed in future work. Firstly, the method was trained on a restrictive dataset derived from the core MIBiG dataset of 2,046 BGCs, which provides limited sequence and product diversity for the model to learn from. To alleviate this issue, we introduced a data augmentation scheme that related additional BGC Atlas sequences to MIBiG clusters. Although these augmented BGC-product class relationships are less reliable than the human-curated MIBiG annotations, we observed that their inclusion in the dataset has been beneficial. It stands to reason that expanding the dataset – whether with additional high-quality curated labels or augmented “noisy” labels – would further enhance the model. Secondly, our method relies on NPClassifier predictions for labeling products in the dataset, which uses a deep neural network that may not always yield the correct results. Unfortunately, there is currently no manually curated dataset that provides detailed classification of BGC products; however, we anticipate that this situation will improve over time. Finally, the ESM Cambrian model was originally trained on protein sequences and as such may not be ideal for embedding BGC genes. Future work may consider the use of fine tuning or end-to-end training of the language model on genomic datasets to improve its match to tasks involving biosynthetic genes.

However, despite these limitations, our technique remains effective for detailed BGC product classification and is the first one to emphasize the importance of using a nomenclature designed for natural products. By using a fine-grained system for describing product types, *BGCat* offers an unmatched insight into the products of individual BGCs and GCFs in aggregate. We expect that this additional context will lead to advances in product structure prediction and, more broadly, an improved understanding of BGC functions.

## 5 Data Availability

The data used in this work is based on publicly available datasets. The MIBiG datasets are available at https://mibig.secondarymetabolites.org. The BGC Atlas dataset can be downloaded from https://bgc-atlas.cs.uni-tuebingen.de. AntiSMASH DB is available at https://antismash-db.secondarymetabolites.org. Our trained model and inference code are available at https://github.com/HassounLab/BGCat.

## References

[1] David J. Newman and Gordon M. Cragg. Natural products as sources of new drugs over the nearly four decades from 01/1981 to 09/2019. Journal of Natural Products, 83(3):770–803, 3 2020.

[2] Arnold L Demain. Importance of microbial natural products and the need to revitalize their discovery. Journal of Industrial Microbiology and Biotechnology, 41(2):185–201, 02 2014.

[3] Manoj Jangra, Dmitrii Y. Travin, Elena V. Aleksandrova, Manpreet Kaur, Lena Darwish, Kalinka Koteva, Dorota Klepacki, Wenliang Wang, Maya Tiffany, Akosiererem Sokaribo, Xuefei Chen, Zixin Deng, Meifeng Tao, Brian K. Coombes, Nora Vazquez-Laslop, Yury S. Polikanov, Alexander S. Mankin, and Gerard D. Wright. A broad-spectrum lasso peptide antibiotic targeting the bacterial ribosome. Nature, 640(8060):1022–1030, Apr 2025.

[4] W. Leimgruber, A. D. Batcho, and F. Schenker. The structure of anthramycin. Journal of the American Chemical Society, 87(24):5793–5795, Dec 1965.

[5] Ben Shen. A new golden age of natural products drug discovery. Cell, 163(6):1297–1300, 12 2015.

[6] Ray Chen, Hon Lun Wong, and Brendan Paul Burns. New approaches to detect biosynthetic gene clusters in the environment. Medicines, 6(1), 2019.

[7] Inge Kjærbølling, Uffe H. Mortensen, Tammi Vesth, and Mikael R. Andersen. Strategies to establish the link between biosynthetic gene clusters and secondary metabolites. Fungal Genetics and Biology, 130:107–121, 2019.

[8] Geoffrey D Hannigan, David Prihoda, Andrej Palicka, Jindrich Soukup, Ondrej Klempir, Lena Rampula, Jindrich Durcak, Michael Wurst, Jakub Kotowski, Dan Chang, Rurun Wang, Grazia Piizzi, Gergely Temesi, Daria J Hazuda, Christopher H Woelk, and Danny A Bitton. A deep learning genome-mining strategy for biosynthetic gene cluster prediction. Nucleic Acids Research, 47(18):e110–e110, 08 2019.

[9] Carolina Rios-Martinez, Nicholas Bhattacharya, Ava P. Amini, Lorin Crawford, and Kevin K. Yang. Deep self-supervised learning for biosynthetic gene cluster detection and product classification. PLOS Computational Biology, 19(5):1–14, 05 2023.

[10] Zhihua Du, Ningyu Zhong, and Jianqiang Li. Enhancing gene cluster identification and classification in bacterial genomes through synonym replacement and deep learning. In 2024 IEEE International Conference on Bioinformatics and Biomedicine (BIBM), pages 19–24, 2024.

[11] Kai Blin, Simon Shaw, Hannah E Augustijn, Zachary L Reitz, Friederike Biermann, Mohammad Alanjary, Artem Fetter, Barbara R Terlouw, William W Metcalf, Eric J N Helfrich, Gilles P van Wezel, Marnix H Medema, and Tilmann Weber. antismash 7.0: new and improved predictions for detection, regulation, chemical structures and visualisation. Nucleic Acids Research, 51(W1):W46–W50, 05 2023.

[12] Kai Blin, Simon Shaw, Lisa Vader, Judit Szenei, Zachary L Reitz, Hannah E Augustijn, José D D Cediel-Becerra, Valérie de Crécy-Lagard, Robert A Koetsier, Sam E Williams, Pablo Cruz-Morales, Sopida Wongwas, Alejandro E Segurado Luchsinger, Friederike Biermann, Aleksandra Korenskaia, Mitja M Zdouc, David Meijer, Barbara R Terlouw, Justin J J van der Hooft, Nadine Ziemert, Eric J N Helfrich, Joleen Masschelein, Christophe Corre, Marc G Chevrette, Gilles P van Wezel, Marnix H Medema, and Tilmann Weber. antismash 8.0: extended gene cluster detection capabilities and analyses of chemistry, enzymology, and regulation. Nucleic Acids Research, 53(W1):W32–W38, 04 2025.

[13] Priyesh Agrawal, Shradha Khater, Money Gupta, Neetu Sain, and Debasisa Mohanty. Rippminer: a bioinformatics resource for deciphering chemical structures of ripps based on prediction of cleavage and cross-links. Nucleic Acids Research, 45(W1):W80–W88, 05 2017.

[14] Eric J. N. Helfrich, Reiko Ueoka, Alon Dolev, Michael Rust, Roy A. Meoded, Agneya Bhushan, Gianmaria Califano, Rodrigo Costa, Muriel Gugger, Christoph Steinbeck, Pablo Moreno, and Jörn Piel. Automated structure prediction of trans-acyltransferase polyketide synthase products. Nature Chemical Biology, 15(8):813–821, 8 2019.

[15] Michael A. Skinnider, Chad W. Johnston, Mathusan Gunabalasingam, Nishanth J. Merwin, Agata M. Kieliszek, Robyn J. MacLellan, Haoxin Li, Michael R. M. Ranieri, Andrew L. H. Webster, My P. T. Cao, Annabelle Pfeifle, Norman Spencer, Q. Huy To, Dan Peter Wallace, Chris A. Dejong, and Nathan A. Magarvey. Comprehensive prediction of secondary metabolite structure and biological activity from microbial genome sequences. Nature Communications, 11(1):6058, 11 2020.

[16] Martin Larralde and Georg Zeller. Machine learning inference of natural product chemistry across biosynthetic gene cluster types. bioRxiv, 2025.

[17] Kai Dührkop, Louis-Félix Nothias, Markus Fleischauer, Raphael Reher, Marcus Ludwig, Martin A. Hoffmann, Daniel Petras, William H. Gerwick, Juho Rousu, Pieter C. Dorrestein, and Sebastian Böcker. Systematic classification of unknown metabolites using high-resolution fragmentation mass spectra. Nature Biotechnology, 39(4):462–471, 4 2021.

[18] Hyun Woo Kim, Mingxun Wang, Christopher A. Leber, Louis-Félix Nothias, Raphael Reher, Kyo Bin Kang, Justin J. J. van der Hooft, Pieter C. Dorrestein, William H. Gerwick, and Garrison W. Cottrell. Npclassifier: A deep neural network-based structural classification tool for natural products. Journal of Natural Products, 84(11):2795–2807, 2021. PMID: 34662515.

[19] Mitja M Zdouc, Kai Blin, Nico L L Louwen, Jorge Navarro, Catarina Loureiro, Chantal D Bader, Constance B Bailey, Lena Barra, Thomas J Booth, Kenan A J Bozhüyük, José D D Cediel-Becerra, Zachary Charlop-Powers, Marc G Chevrette, Yit Heng Chooi, Paul M D’Agostino, Tristan de Rond, Elena Del Pup, Katherine R Duncan, Wenjia Gu, Novriyandi Hanif, Eric J N Helfrich, Matthew Jenner, Yohei Katsuyama, Aleksandra Korenskaia, Daniel Krug, Vincent Libis, George A Lund, Shrikant Mantri, Kalindi D Morgan, Charlotte Owen, Chin-Soon Phan, Benjamin Philmus, Zachary L Reitz, Serina L Robinson, Kumar Saurabh Singh, Robin Teufel, Yaojun Tong, Fidele Tugizimana, Dana Ulanova, Jaclyn M Winter, César Aguilar, Daniel Y Akiyama, Suhad A A Al-Salihi, Mohammad Alanjary, Fabrizio Alberti, Gajender Aleti, Shumukh A Alharthi, Mariela Y Arias Rojo, Amr A Arishi, Hannah E Augustijn, Nicole E Avalon, J Abraham Avelar-Rivas, Kyle K Axt, Hellen B Barbieri, Julio Cesar J Barbosa, Lucas Gabriel Barboza Segato, Susanna E Barrett, Martin Baunach, Christine Beemelmanns, Dardan Beqaj, Tim Berger, Jordan Bernaldo-Agüero, Sandra M Bettenbühl, Vincent A Bielinski, Friederike Biermann, Ricardo M Borges, Rainer Borriss, Milena Breitenbach, Kevin M Bretscher, Michael W Brigham, Larissa Buedenbender, Brodie W Bulcock, Carolina Cano-Prieto, Joao Capela, Victor J Carrion, Riley S Carter, Raquel Castelo-Branco, Gabriel Castro-Falcon, Fernanda O Chagas, Esteban Charria-Girón, Ayesha Ahmed Chaudhri, Vasvi Chaudhry, Hyukjae Choi, Yukyung Choi, Roya Choupannejad, Jakub Chromy, Melinda S Chue Donahey, Jérôme Collemare, Jack A Connolly, Kaitlin E Creamer, Max Crüsemann, Andres Arredondo Cruz, Andres Cumsille, Jean-Felix Dallery, Luis Caleb Damas-Ramos, Tito Damiani, Martinus de Kruijff, Belén Delgado Martín, Gerardo Della Sala, Jelle Dillen, Drew T Doering, Shravan R Dommaraju, Suhan Durusu, Susan Egbert, Mark Ellerhorst, Baptiste Faussurier, Artem Fetter, Marc Feuermann, David P Fewer, Jonathan Foldi, Andri Frediansyah, Erin A Garza, Athina Gavriilidou, Andrea Gentile, Jennifer Gerke, Hans Gerstmans, Juan Pablo Gomez-Escribano, Luz A González-Salazar, Natalie E Grayson, Claudio Greco, Juan E Gris Gomez, Sebastian Guerra, Shaday Guerrero Flores, Alexey Gurevich, Karina Gutiérrez-García, Lauren Hart, Kristina Haslinger, Beibei He, Teo Hebra, Jethro L Hemmann, Hindra Hindra, Lars Höing, Darren C Holland, Jonathan E Holme, Therese Horch, Pavlo Hrab, Jie Hu, Thanh-Hau Huynh, Ji-Yeon Hwang, Riccardo Iacovelli, Dumitrita Iftime, Marianna Iorio, Sidharth Jayachandran, Eunah Jeong, Jiayi Jing, Jung J Jung, Yuya Kakumu, Edward Kalkreuter, Kyo Bin Kang, Sangwook Kang, Wonyong Kim, Geum Jin Kim, Hyunwoo Kim, Hyun Uk Kim, Martin Klapper, Robert A Koetsier, Cassandra Kollten, Ákos T Kovács, Yelyzaveta Kriukova, Noel Kubach, Aditya M Kunjapur, Aleksandra K Kushnareva, Andreja Kust, Jessica Lamber, Martin Larralde, Niels J Larsen, Adrien P Launay, Ngoc-Thao-Hien Le, Sarah Lebeer, Byung Tae Lee, Kyungha Lee, Katherine L Lev, Shu-Ming Li, Yong-Xin Li, Cuauhtémoc Licona-Cassani, Annette Lien, Jing Liu, Julius Adam V Lopez, Nataliia V Machushynets, Marla I Macias, Taifo Mahmud, Matiss Maleckis, Añadir Maharai Martinez-Martinez, Yvonne Mast, Marina F Maximo, Christina M McBride, Rose M McLellan, Khyati Mehta Bhatt, Chrats Melkonian, Aske Merrild, Mikko Metsä-Ketelä, Douglas A Mitchell, Alison V Müller, Giang-Son Nguyen, Hera T Nguyen, Timo H J Niedermeyer, Julia H O’Hare, Adam Ossowicki, Bohdan O Ostash, Hiroshi Otani, Leo Padva, Sunaina Paliyal, Xinya Pan, Mohit Panghal, Dana S Parade, Jiyoon Park, Jonathan Parra, Marcos Pedraza Rubio, Huong T Pham, Sacha J Pidot, Jörn Piel, Bita Pourmohsenin, Malik Rakhmanov, Sangeetha Ramesh, Michelle H Rasmussen, Adriana Rego, Raphael Reher, Andrew J Rice, Augustin Rigolet, Adriana Romero-Otero, Luis Rodrigo Rosas-Becerra, Pablo Y Rosiles, Adriano Rutz, Byeol Ryu, Libby-Ann Sahadeo, Murrel Saldanha, Luca Salvi, Eduardo Sánchez-Carvajal, Christian Santos-Medellin, Nicolau Sbaraini, Sydney M Schoellhorn, Clemens Schumm, Ludek Sehnal, Nelly Selem, Anjali D Shah, Tania K Shishido, Simon Sieber, Velina Silviani, Garima Singh, Hemant Singh, Nika Sokolova, Eva C Sonnenschein, Margherita Sosio, Sven T Sowa, Karin Steffen, Evi Stegmann, Alena B Streiff, Alena Strüder, Frank Surup, Tiziana Svenningsen, Douglas Sweeney, Judit Szenei, Azat Tagirdzhanov, Bin Tan, Matthew J Tarnowski, Barbara R Terlouw, Thomas Rey, Nicola U Thome, Laura Rosina Torres Ortega, Thomas Tørring, Marla Trindade, Andrew W Truman, Marie Tvilum, Daniel W Udwary, Christoph Ulbricht, Lisa Vader, Gilles P van Wezel, Max Walmsley, Randika Warnasinghe, Heiner G Weddeling, Angus N M Weir, Katherine Williams, Sam E Williams, Thomas E Witte, Steffaney M Wood Rocca, Keith Yamada, Dong Yang, Dongsoo Yang, Jingwei Yu, Zhenyi Zhou, Nadine Ziemert, Lukas Zimmer, Alina Zimmermann, Christian Zimmermann, Justin J J van der Hooft, Roger G Linington, Tilmann Weber, and Marnix H Medema. Mibig 4.0: advancing biosynthetic gene cluster curation through global collaboration. Nucleic Acids Research, 53(D1):D678–D690, 12 2024.

[20] Satria A Kautsar, Kai Blin, Simon Shaw, Tilmann Weber, and Marnix H Medema. Big-fam: the biosynthetic gene cluster families database. Nucleic Acids Research, 49(D1):D490–D497, 10 2020.

[21] Satria A Kautsar, Justin J J van der Hooft, Dick de Ridder, and Marnix H Medema. Big-slice: A highly scalable tool maps the diversity of 1.2 million biosynthetic gene clusters. GigaScience, 10(1):giaa154, 01 2021.

[22] Peter Cimermancic, Marnix H. Medema, Jan Claesen, Kenji Kurita, Laura C. Wieland Brown, Konstantinos Mavrommatis, Amrita Pati, Paul A. Godfrey, Michael Koehrsen, Jon Clardy, Bruce W. Birren, Eriko Takano, Andrej Sali, Roger G. Linington, and Michael A. Fischbach. Insights into secondary metabolism from a global analysis of prokaryotic biosynthetic gene clusters. Cell, 158(2):412–421, 7 2014.

[23] James R. Doroghazi, Jessica C. Albright, Anthony W. Goering, Kou-San Ju, Robert R. Haines, Konstantin A. Tchalukov, David P. Labeda, Neil L. Kelleher, and William W. Metcalf. A roadmap for natural product discovery based on large-scale genomics and metabolomics. Nature Chemical Biology, 10(11):963– 968, 11 2014.

[24] Tiago Leao, Guilherme Castelao, Anton Korobeynikov, Emily A. Monroe, Sheila Podell, Evgenia Glukhov, Eric E. Allen, William H. Gerwick, and Lena Gerwick. Comparative genomics uncovers the prolific and distinctive metabolic potential of the cyanobacterial genus ¡i¿moorea¡/i¿. Proceedings of the National Academy of Sciences, 114(12):3198–3203, 2017.

[25] Jorge C. Navarro-Muñoz, Nelly Selem-Mojica, Michael W. Mullowney, Satria A. Kautsar, James H. Tryon, Elizabeth I. Parkinson, Emmanuel L. C. De Los Santos, Marley Yeong, Pablo Cruz-Morales, Sahar Abubucker, Arne Roeters, Wouter Lokhorst, Antonio Fernandez-Guerra, Luciana Teresa Dias Cappelini, Anthony W. Goering, Regan J. Thomson, William W. Metcalf, Neil L. Kelleher, Francisco Barona-Gomez, and Marnix H. Medema. A computational framework to explore large-scale biosynthetic diversity. Nature Chemical Biology, 16(1):60–68, 1 2020.

[26] Anthony W. Goering, Ryan A. McClure, James R. Doroghazi, Jessica C. Albright, Nicole A. Haverland, Yongbo Zhang, Kou-San Ju, Regan J. Thomson, William W. Metcalf, and Neil L. Kelleher. Metabologenomics: Correlation of microbial gene clusters with metabolites drives discovery of a nonribosomal peptide with an unusual amino acid monomer. ACS Central Science, 2(2):99–108, 2 2016.

[27] ESM Team. Esm cambrian: Revealing the mysteries of proteins with unsupervised learning, 2024.

[28] Kai Blin, Simon Shaw, Marnix H Medema, and Tilmann Weber. The antismash database version 4: additional genomes and bgcs, new sequence-based searches and more. Nucleic Acids Research, 52(D1):D586–D589, 10 2023.

[29] RDKit contributors. RDKit: Open-source cheminformatics. Release 2022.09.1, 10 2022.

[30] Diederik P. Kingma and Jimmy Ba. Adam: A method for stochastic optimization, 2017.

[31] Caner Bagcı, Matin Nuhamunada, Hemant Goyat, Casimir Ladanyi, Ludek Sehnal, Kai Blin, Satria A Kautsar, Azat Tagirdzhanov, Alexey Gurevich, Shrikant Mantri, Christian von Mering, Daniel Udwary, Marnix H Medema, Tilmann Weber, and Nadine Ziemert. Bgc atlas: a web resource for exploring the global chemical diversity encoded in bacterial genomes. Nucleic Acids Research, 53(D1):D618–D624, 10 2024.

[32] Lorna Richardson, Ben Allen, Germana Baldi, Martin Beracochea, Maxwell L Bileschi, Tony Burdett, Josephine Burgin, Juan Caballero-Pérez, Guy Cochrane, Lucy J Colwell, Tom Curtis, Alejandra Escobar-Zepeda, Tatiana A Gurbich, Varsha Kale, Anton Korobeynikov, Shriya Raj, Alexander B Rogers, Ekaterina Sakharova, Santiago Sanchez, Darren J Wilkinson, and Robert D Finn. Mgnify: the microbiome sequence data analysis resource in 2023. Nucleic Acids Research, 51(D1):D753–D759, 12 2022.

[33] Elizabeth A. Freeman and Gretchen G. Moisen. A comparison of the performance of threshold criteria for binary classification in terms of predicted prevalence and kappa. Ecological Modelling, 217(1):48–58, 2008.

[34] Charles Parker. An analysis of performance measures for binary classifiers. In 2011 IEEE 11th International Conference on Data Mining, pages 517–526, 2011.

[35] Eric J. Martin, Valery R. Polyakov, Li Tian, and Rolando C. Perez. Profile-qsar 2.0: Kinase virtual screening accuracy comparable to four-concentration ic50s for realistically novel compounds. Journal of Chemical Information and Modeling, 57(8):2077–2088, 2017. PMID: 28651433.

[36] Gurjeet S Kohli, Uwe John, Frances M Van Dolah, and Shauna A Murray. Evolutionary distinctiveness of fatty acid and polyketide synthesis in eukaryotes. The ISME Journal, 10(8):1877–1890, 01 2016.

[37] Justin J. J. van der Hooft, Hosein Mohimani, Anelize Bauermeister, Pieter C. Dorrestein, Katherine R. Duncan, and Marnix H. Medema. Linking genomics and metabolomics to chart specialized metabolic diversity. Chem. Soc. Rev., 49:3297–3314, 2020.

[38] Satria A. Kautsar, Hernando G. Suarez Duran, Kai Blin, Anne Osbourn, and Marnix H. Medema. plantismash: automated identification, annotation and expression analysis of plant biosynthetic gene clusters. Nucleic Acids Research, 45(W1):W55–W63, 04 2017.

[39] Elena Del Pup, Charlotte Owen, Ziqiang Luo, Hannah E. Augustijn, Arjan Draisma, Guy Polturak, Satria A. Kautsar, Anne Osbourn, Justin J.J. van der Hooft, and Marnix H. Medema. plantismash 2.0: improvements to detection, annotation, and prioritization of plant biosynthetic gene clusters. bioRxiv, 2025.

[40] Taehyung Kwon and Blake T. Hovde. Global characterization of biosynthetic gene clusters in non-model eukaryotes using domain architectures. Scientific Reports, 14(1):1534, 1 2024.

