## Supplementary Materials for "Fine-Grained Structural Classification of Biosynthetic Gene Cluster-Encoded Products"

This section lists all supplementary files referred to in the main text and included in the zip archive.

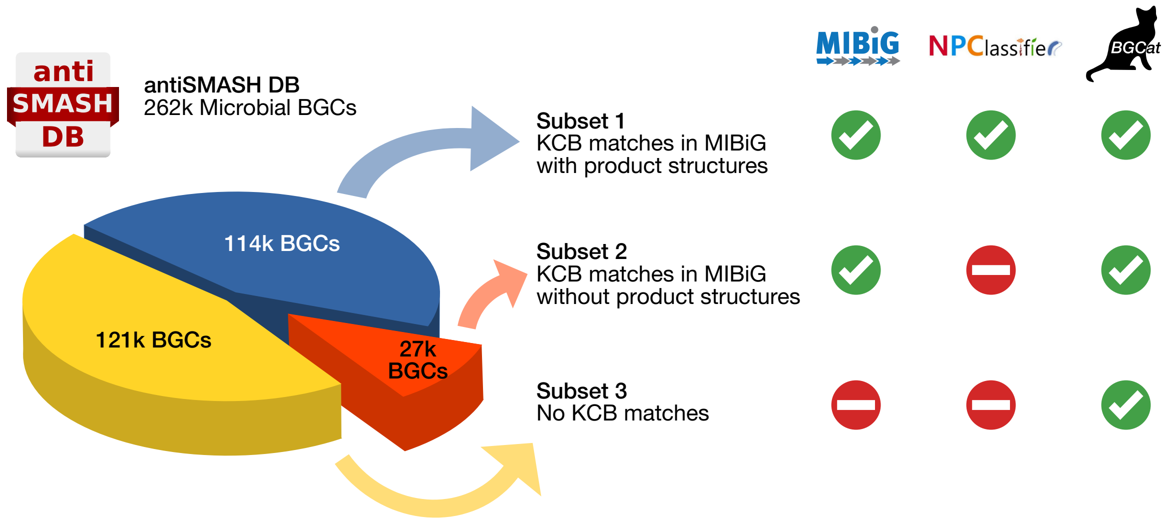


**Supplementary Figure 1:** An overview of the three antiSMASH DB subsets. BGCs from subset 1 feature KCB matches to MIBiG BGCs with product structures, allowing the use of MIBiG labels and NPClassifier predictions for BGC characterization. Subset 2 also has matches to MIBiG BGCs, albeit without machine-readable product structures, precluding the use of NPClassifier. Finally, subset 3 features no KCB matches at all, which eliminates the availability of MIBiG labels as well. However, *BGCat* remain available for all three subsets.

**Supplementary Table 1.** List of BGC Atlas BGCs and their corresponding GCFs. Predicted product class labels are also included for each BGC.

**Supplementary Table 2.** List of GCFs generated based on BGC Atlas, indicating whether or not a given family has a distinct PCP. In addition, labels for the most overrepresented product classes are included for each GCF.

**Supplementary Table 3.** List of antiSMASH DB BGCs from subset 3 that lack KCB hits in MIBiG. For each BGC, we provide the top 3 *BGCat* product classification predictions.
